# scAEGAN: Unification of Single-Cell Genomics Data by Adversarial Learning of Latent Space Correspondences

**DOI:** 10.1101/2022.04.19.488745

**Authors:** Sumeer Ahmad Khan, Robert Lehmann, Xabier Martinez-de-Morentin, Albert Malillo Ruiz, Vincenzo Lagani, Narsis A. Kiani, David Gomez-Cabrero, Jesper Tegner

## Abstract

Recent progress in Single-Cell Genomics have produced different library protocols and techniques for profiling of one or more data modalities in individual cells. Machine learning methods have separately addressed specific integration challenges (libraries, samples, paired-unpaired data modalities). We formulate an unifying data-driven methodology addressing all these challenges. To this end, we design a hybrid architecture using an autoencoder (AE) network together with adversarial learning by a cycleGAN (cGAN) network, jointly referred to as scAEGAN. The AE learns a low-dimensional embedding of each condition, whereas the cGAN learns a non-linear mapping between the AE representations. The core insight is that the AE respects each sample’s uniqueness, whereas the cGAN exploits the distributional data similarity in the latent space. We evaluate scAEGAN using simulated data and real datasets of a single-modality (scRNA-seq), different library preparations (Fluidigm C1, CelSeq, CelSeq2, SmartSeq), and several data modalities such as paired scRNA-seq and scATAC-seq. We find that scAEGAN outperforms Seurat3 in library integration, is more robust against data sparsity, and beats Seurat 4 in integrating paired data from the same cell. Furthermore, in predicting one data modality from another, scAEGAN outperforms Babel. We conclude scAEGAN surpasses current state-of-the-art methods across several seemingly different integration challenges.

## INTRODUCTION

The maturation of the single-cell genomics field has produced methods able to profile multiple data modalities, such as single-cell RNA sequencing (scRNA-seq) and chromatin profiles (scATAC-seq), even on the same cells at the same time. This development has provided rich opportunities for a deep understanding of cell states and transitions while presenting severe computational challenges (Stuart et al., 2019). One of the most notable challenges is the integration of different single-cell datasets. Integrating different experiments has proved daunting even when using the same library protocol and omics type. For example, distinct scRNA-seq datasets may differ in the number of sampled cells and sequencing depth allocated to each cell, even by several orders of magnitude. The next challenge is to combine scRNA-seq data originating from different library protocols or species (Shafer, 2019). A third challenging task is integrating other data modalities from the same experiment but originating from separate cells, a case known as unpaired multi-omics integration. Finally, recent technological advances produce paired multi-omics data collected from the same cell. These challenges have thus far been addressed one by one. For example, Seurat3 (Stuart & Satija, 2019) and MOFA+ (Argelaguet et al., 2020) integrate unpaired data, whereas Seurat4 (Hao et al., 2021) integrates paired data and Babel (Wu et al., 2021) predicts one modality from another. Methods such as scAlign (Johansen & Quon, 2019), Harmony (Korsunsky et al., 2019), and Seurat3 target scRNA-seq datasets originating from different experiments that used the same platform (Tran et al., 2020).

In contrast, Liger (Welch et al., 2019), scMerge (Lin et al., 2019), and Seurat3 can integrate datasets produced using different library protocols. Most of these limitations directly derive from the internal operation of each method. Seurat3 is based on the concept of “anchors”, which are cross-datasets pairs of cells that share a similar biological state. This approach does not readily scale to large datasets and performs poorly when integrating heterogeneous datasets (Li et al., 2022). Worse, only a fraction of cell types is usually shared across datasets, making identification increasingly challenging using anchors (Y. Zhang & Wang, 2021). Babel, a machine learning method, targets only gene prediction for paired data. Thus, by design, it lacks clustering capabilities and cannot tackle unpaired data or different library protocols.

Furthermore, these approaches implicitly assume that differences between datasets arise entirely from technical variation, thus potentially masking the biological signal. For example, the Mutual Nearest Neighbors (MNNs) method (Haghverdi et al., 2018) effectively reduces differences between datasets. An alternative strategy is exemplified by Seurat3 (Welch et al., 2019), which forces all datasets into a shared latent space. However, both dataset similarities and differences in many kinds of analysis are biologically meaningful. Thus, it requires respecting each sample’s uniqueness, protocol, and data type.

There is a need for scalable and robust integrative methods for omics data, preferentially general enough to encompass the multitude of integration tasks in one systematic framework. Not the least, from this standpoint of that, we can expect the scale and the number of different data modalities to further increase in the future.

Here we present a novel integrative method that complies with all such requirements. The key point of our approach is that we do not force all experimental samples into a single joint representation, regardless of their library protocol, data modality, paired or unpaired design. Instead, we use an autoencoder (AE) to represent and respect the distributional characteristics of each dataset and condition. The integration is performed in the latent space by learning a mapping between the different latent space representations. Inspired by recent progress in image-to-image translation, we use a cycleGAN (cGAN) architecture for obtaining a translation between the latent spaces corresponding to different datasets. We denote our method scAEGAN, a coupled AE – cycleGAN architecture. Our results demonstrate that scAEGAN can target single-cell multi-omics integration tasks with performances similar to or superior to other state-of-the-art tools. Furthermore, we provide evidence that the mapping between different latent spaces is essential for effective integration by contrasting scAEGAN against the simplified approach of directly concatenating latent spaces, which forces the data into a shared latent space without learning a mapping.

## MATERIAL AND METHODS

### Neural Network Architecture

scAEGAN is a unifying architecture combining AE (Hinton & Salakhutdinov, 2006) and cGAN (Zhu et al., 2017). AE, an unsupervised deep neural network, learns essential latent features and ignores the non-essential sources of variations such as random noise (Eraslan et al., 2019). Hence the high dimensional ambient space is represented in a compressed form, capturing the underlying proper data manifold.

First, each given dataset is provided as input to an AE in a matrix *X*, where rows (m) represent the cells and columns (n) indicate genes/transcripts. The AE task consists in learning the encoding representation through an encoding function *e*(*x*) and then mapping back *e*(*x*) to the original input space through a decoding function *d*. For faster convergence and better accuracy, Rectified Linear Unit (ReLU) has been used as an activation function, which is given as a function *f* applied to the input *x* :

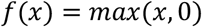

The first hidden layer *Hidden*_1_ with *l*_1_ nodes following the input *X*_*i*_ row vector) is formulated as follows:

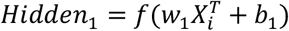

The weight matrix *w*_1_ is of *l*_1_ × *n* dimensions and the bias term *b*_1_ is *l*_1_ length vector. Each subsequent middle layer *k* is formulated as:

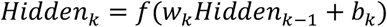

The composition of *e* and *d*, i.e., *d*(*e*(*x*)) = *X*^′^ is called the reconstruction function, and the reconstruction loss function penalizes the error made, which is given as:

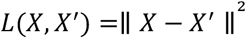

The low dimensional space representation from the AEs captures the underlying manifold of the data. Secondly, we utilize a cGAN to learn relationships between the different domains/datasets (A and B). Specifically, learning two generative mapping functions *G*_*AB*_ : *A*−> *B* and *G*_*BA*_ : *B*−> *A*. In addition to these generative functions, two discriminators *D*_*A*_ and *D*_*B*_ were used to regularise the generators to generate samples from a distribution close to the latent representation of A or B. We used the Wasserstein GAN adversarial loss introduced in (Arjovsky et al., 2017). In the Wasserstein GAN, the discriminator is replaced by a critic model. The function of the critic is not directly telling the fake samples apart from the real ones. Instead, it is trained to learn a *K*-Lipschitz continuous function, which makes the neural network gradient smaller than a threshold value *K*, such that ‖∇*f*‖ ≤ *K*. The primary rationale for applying this condition is that gradient behaves better, making generator optimization easier(Qin et al., 2018). As the loss function decreases in training, the Wasserstein distance gets smaller, and the generator model’s output grows closer to the actual data distribution. This Wasserstein GAN adversarial loss is applied to both the mapping functions, and the objective is expressed for *G*_*AB*_ : *A*−> *B*:

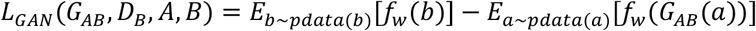

where function *f* is a K-Lipschitz continuous function, {*f*_*w*_}_*w* ∈*W*_, parameterized by *w*. The cycle consistency loss ensures that the learned mappings are cycle consistent, i.e., be able to bring back to the original domain. Which is given as:

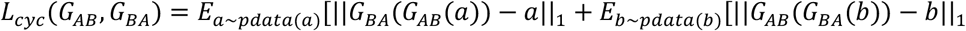

To train the cGAN on the latent subspaces of the two domains, the entire objective function is:

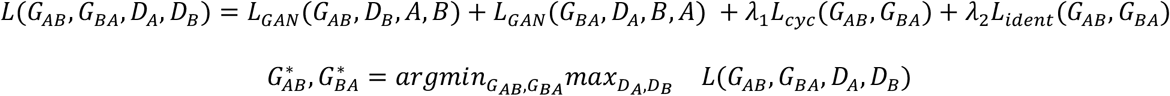

The scAEGAN architecture is provided with a scRNA-seq and a scATAC-seq data set (domain A and B, respectively) as illustrated in Figure 1a. The first step in the scAEGAN integration algorithm is training an AE independently on both domains A and B to find a low-dimensional embedding that preserves each domain’s key defining features. This step is necessary since direct translation between scRNA-seq domains via cGAN, while possible, is hampered by increased technical variation or dataset complexity. AE is particularly suitable due to their ability to reduce random noise while still maintaining essential features. Moreover, it turns out that AE generates more biologically meaningful embeddings than variational autoencoders (VAE) when learning across latent spaces. A cGAN is then trained on the low-dimensional representations to achieve the translation between domains.

**Figure. 1.**
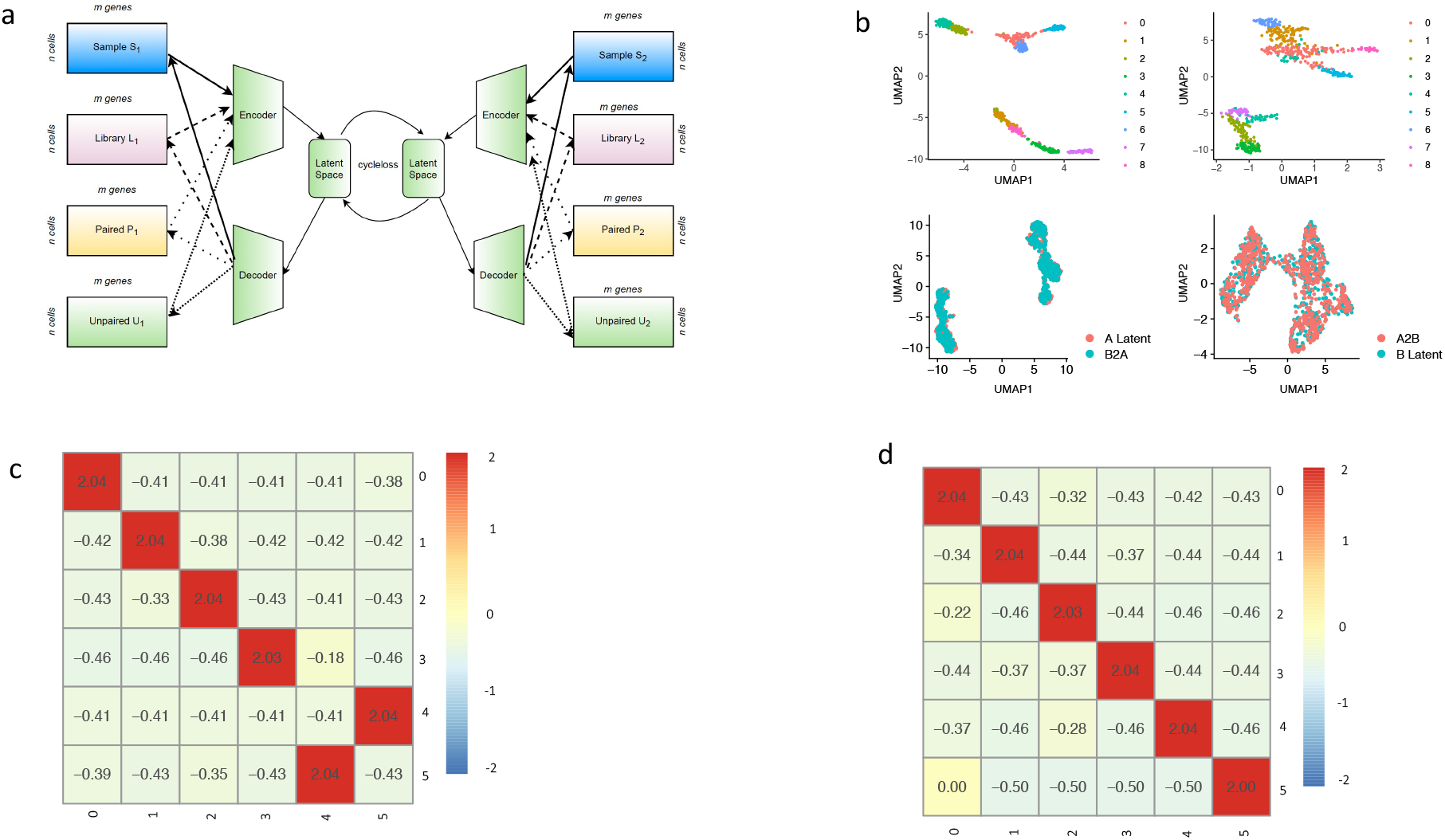
scAEGAN architecture for single cell data integration **a)** Coupled scAEGAN, allowing the translation of AE-obtained low-dimensional embeddings via a cGAN, **b)** Outputs from the scAEGAN, where A2B and B2A are integration results of the A and B datasets, **c, d)** scAEGAN preserved the transferred cell identity agreement with the original identity.

### Hyperparameter Tuning

We have performed a series of analyses to generate the best configuration for scAEGAN hyperparameters (layers, the number of nodes, the activation function used, and the optimizer used) based on the nature of the single-cell data, that resulted in the optimal configuration.

### AE Hyperparameter and optimization

The AE model consists of three hidden layers with the dimensions of *Hidden*_1_(300), *Hidden*_2_(50), *Hidden*_3_(300). ReLU has been used as an activation function followed by a Sigmoid activation function in the output layer. A dropout value of 0.2 has been used to prevent overfitting. We used Adam (Kingma & Ba, 2015) as an optimizer with different settings ranging from lr= 0.0001 to 0.0005 and found the 0.0001 as the best setting for our experiments to train the AE model. We trained the AE using a batch size of 16 for the number of epochs ranging from 60 to 250 and observed that the model trained for 120 epochs gives better performance, with 80% training and 20% validation data to analyze the convergence of the model.

### cGAN Hyperparameter and optimization

The architecture of cGAN consists of two generators and two discriminators. The generators consist of one residual block and one dense layer of 50 dimensions each, and the discriminators consist of two dense layers. For residual block, a dropout value of (0.2) has been used, followed by batch normalization, which stabilizes the learning process. In addition to this, batch normalization has a slight regularization effect; for this reason, we have used a small value (0.2) for the dropout. LeakyReLU(Maas et al., 2013) has been used as an activation function. To train the cGAN model, we have used two different optimized settings of Adam optimizer for the real data and simulated data. For the actual data, the cGAN is trained with Adam optimizer with parameters: *lr* = 0.0005 *and r* = 0.0002, *beta*_1_ = 0.5, *beta*_2_ = 0.999, *epsilon* = 1*e* − 7, *decay* = 0. And for the simulated data, the cGAN is trained with Adam optimizer with parameters: *lr* = 0.0002, *beta*_1_ = 0.5, *beta*_2_ = 0.999, *epsilon* = 1*e* − 6, *decay* = 0.0. In all our experiments, we use the hyperparameters as weights for the cyclic loss and identity loss, i.e., λ_1_ = 0.3 and λ_2_ = 0.3, which were chosen to check a couple of combinations for verifying that our optimization process generates the translated data similar to the starting ones. Also, to maintain the K-Lipschitz continuity of *f*_*w*_ we used the hyperparameter *c* = 0.1 during the training, which helps in resulting in compact parameter space. In addition to these, the cGAN is trained with a batch size of 4 for 200 to 400 epochs.

### AE Concatenated (AE-Concat)

The AE concatenated architecture is used for the comparison with scAEGAN. The AE-Concat architecture consists of two encoders of one hidden layer, concatenated and projected down to the bottleneck layer. The first encoder takes the input of the first domain, and the second encoder takes input from the second domain. The first encoder and second encoder dimensions are 30 each, summing up to 60 dimensions after concatenating, projected down to a low dimensional space of 50 dimensions in the bottleneck layer. This layer contains the integrated low-dimensional representation of the two domains. ReLU is used as an activation function and a dropout value of (0.2). This concatenated network is trained with Adam optimizer with a learning rate of *lr* = 0.0005 for 200 epochs using a batch size of 16. The concatenated AE uses the mean square error as a loss function to minimize the input and output loss.

### Overview of the Evaluation Metrics

Firstly, the overlap between datasets before and after integration was visually assessed in low-dimensional representations using the UMAP R package v0.2.3.1. In the case of scAEGAN, integration quality was measured by transferring labels between domains. A support vector machine is first trained to classify cell types in one domain using the cluster assignments obtained from Louvain clustering as implemented in Seurat3. This step is followed by the prediction of cell type in the other domain and a comparison with the original clustering in this domain. In the case of AE integration, direct label transfer between input space and low-dimensional representation of the integrated dataset is not applicable. Accordingly, cell types are again assigned to input and integrated datasets via clustering with the Louvain algorithm and are then directly compared.

Furthermore, Seurat was used to transfer labels using its TransferData function. Cell type assignments, i.e., clusterings, are compared using the Adjusted Rand Index (ARI) in R package pdfCluster v1.0.3. and the Jaccard Index (JI) in R package clusteval v0. In addition to ARI and JI, we used Preserved Pairwise Jaccard Index (PPJI), a non-symmetric distance metric between two clusterings, for evaluating the clustering results.

Since Seurat is the most widely used tool, we compare our integration results with Seurat version 3 and 4 for the different library protocols on paired/unpaired data.

### Jaccard Index (JI)

The JI calculates a 2 by 2 contingency table of agreements and disagreements between the two finite subsets and evaluates the stability of clustering. Given two subsets Ai and Bj, the JI is computed as:

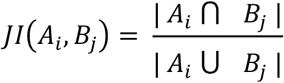

### Adjusted Rand Index (ARI)

The ARI measures the similarity between the two partitions of the same datasets by the proportion of the agreement between the two partitions. The metric is adjusted for chance, such that the independent have an expected index of zero and identical partitions have an ARI equal to 1. The ARI is computed as:

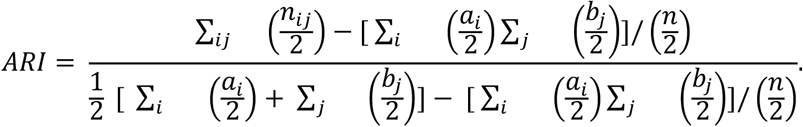

Where, *n*_*ij*_ is the number of common cells between two partitions and *a*_*i*_ = ∑_*k*_ (*n*_*ik*_), *b*_*j*_ = ∑_*k*_ *n*_*jk*_ are the number of cells in estimated cluster i and in true cluster j respectively.

### PredRNA

RNA prediction was carried out by training the cGAN on scRNAseq/scATACseq paired dataset and predicting on the held-out set.

Evaluation for quality check was performed by computing Pearson correlation between each pair of cells from predicted RNA and original RNA training input data. This computation was performed using cor function from stats package.

### Clustering for integrated and independent omic modalities

The Seurat Louvain clustering implementation was used for all of the clustering analysis (Waltman & Van Eck, 2013). Various inputs are considered depending on the analysis:

Single-cell RNA-seq data: PCA components.

Single-cell ATAC-seq data: LSI components.

Cells were clustered based on shared components generated by the methods studied (scAEGAN, Seurat3, Seurat4).

For integrated subspaces, the Louvain resolution has been set to the default value of 0.6. The number of nearest neighbors has been used as K=20.

### Data

For developing and testing this computational approach’s performance and quality, four different datasets (same/different modality, library preparation protocols) have been used. The summary of datasets used is given in Table 1.

**Table 1.**
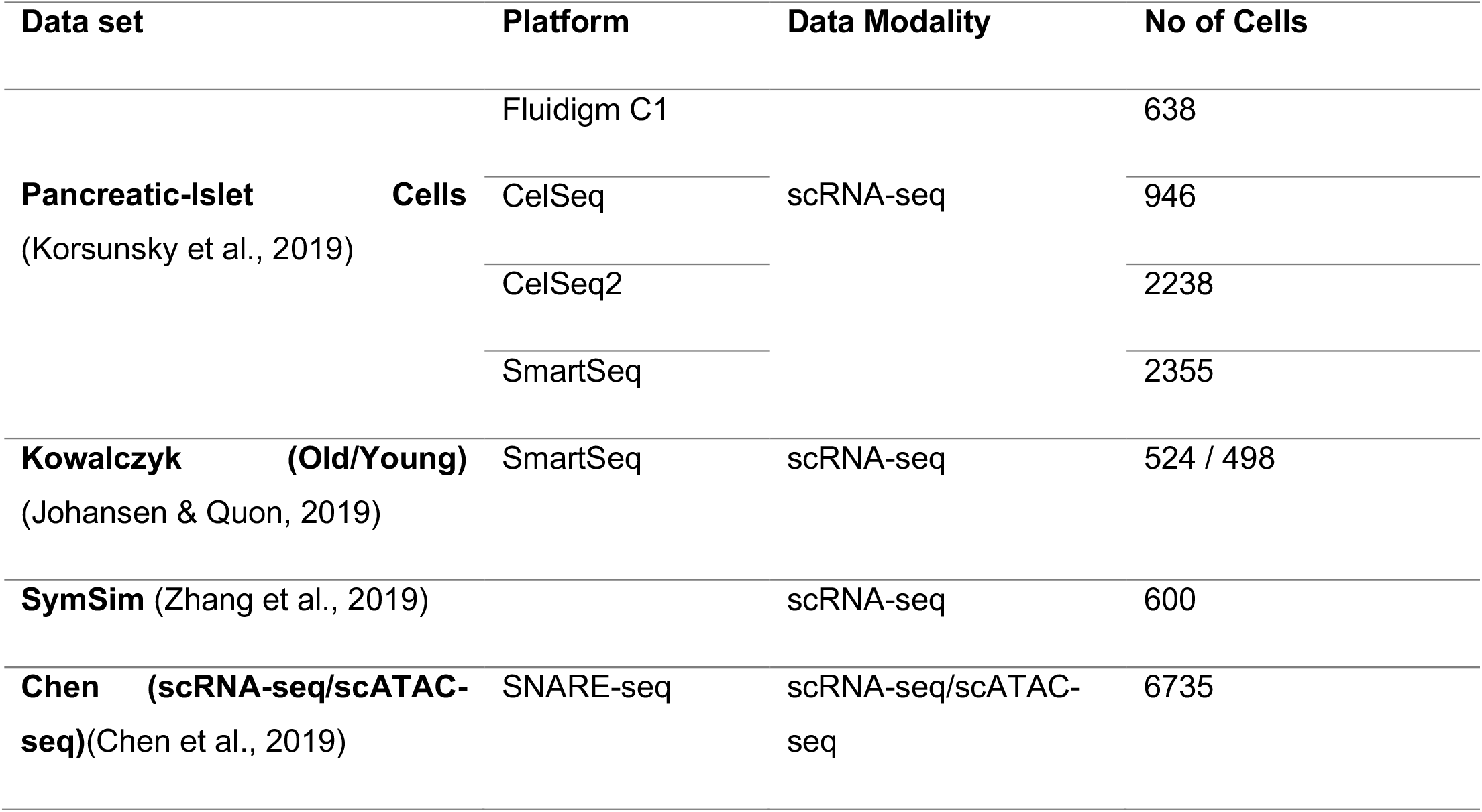
Dataset summary providing data modality, sequencing platform, and number of cells employed for integration after pre-processing.

### Simulated Datasets

Two datasets containing 600 cells from 5 populations and with 3000 genes each were simulated using SymSim (X. Zhang et al., 2019) with the ‘Phyla5’ tree and the following parameters: nevf 35, evf_type ‘continuous’, n_de_evf 5, sigma 0.5, gene_effect_prob 0.5, gene_effect_sd 0.2, alpha_mean 0.05, alpha_sd 0.02, depth_mean 5e4, depth_sd 3e3. For one of these datasets, branch lengths of the ‘Phyla5’ tree was slightly modified.

Two more datasets were simulated for analysis to examine the sufficiency of scAEGAN when there is a cell type unbalance in two datasets. For dataset A, we simulated multiple versions with all cells, 100, 50, and 10 cells for the largest cluster, and for dataset B, we opted to remove the largest cluster, which had about 200 of the 600 genes in it.

### Real Datasets

The pre-processed mouse hematopoietic stem cell dataset of young and old individuals presented by was downloaded from https://github.com/quon-titative-biology/scalign (Johansen & Quon, 2019).

Four human pancreatic islet cell datasets sequenced using different platforms were obtained pre-processed as described in from https://github.com/immunogenomics/harmony2019 (Korsunsky et al., 2019). Raw read count matrices were scaled and normalized using Seurat v3 prior to integration.

scRNAseq/scATACseq paired dataset: We selected an existing paired scRNA-scATAC dataset from the SNARE-seq protocol (a droplet-based single nucleus over mRNA expression and chromatin accessibility sequencing) (Chen et al., 2019). The data was downloaded from GSE126074. The preprocessing applied to this dataset is as follows:

Quality filter – low-quality features: removes low-quality features and cells from both modalities. All cells with an overall abundance level of “number of features per cell” and “number of counts per cell” less than quantile 0.1 and greater than quantile 0.9 were excluded from further analysis. For mRNA (ATAC), minimum abundance filtering was used: genes (peaks) profiled in less than 4 cells (3 cells) and cells with fewer than 201 genes quantified were filtered. There was no requirement for a certain number of peaks per cell. Following quality and abundance filtering, we considered a total of 8,086 cells for scRNA and 8,214 cells for scATAC adult samples for analysis.

ATAC-derived gene activity: To compute ATAC-derived gene activity, the Seurat3 ‘CreateGeneActivityMatrix’ function with “upstream=2000” bases was used. In addition, the GRCh38 genome was used as a reference to later identify marker genes across the integrated expression subspace.

Quality filter – mitochondrial: 5 percent mitochondrial filtering was used for the expression matrices of scRNA and scATAC, with activity from peaks used in the ATAC case.

Component parameters: For scRNA reduction, 15 principal components (PCA) were chosen, and 50 latent semantic indexing components (LSI) were chosen for scATAC.

The final number of cells: from the resulting pipeline, a total of 6,735 paired cell profiles were considered for the downstream analysis.

Integration: On this dataset, Seurat 3 (unpaired) and Seurat 4 (paired) were used to generate a reference integrated version for further processing and later integration. Using standard normalization and integration guides for Seurat3 and Seurat4 Weighted Nearest Neighbor Analysis vignettes (Hao et al., 2021). In Seurat 3, the FindTransferAnchors function was used to generate anchorsets using RNA as the reference and ATAC as the query modalities, with CCA as the reduction method. This was followed by the TransferData function, where the anchorset generated was used to transfer the RNA derived information into the ATAC modality using LSI dimensional reduction for the weighting anchors. Seurat 4 was used to identify multimodal neighbors using the FindMultimodalNeighbors function.

## RESULTS and DISCUSSION

### A novel architectural design for single-cell multi-data set analysis

We propose an integrated AE and cGAN architecture (Fig. 1a), allowing the integration of scRNAseq data from different datasets. A particular experiment in a given data domain produces a cell count matrix, which is then fed into the encoder of the AE to condense it into a lower-dimensional latent representation. The objective of the decoder is to reconstruct the input from the latent representation. This defines a reconstruction loss function for the AE (for details and hyperparameters, see Material and Methods). This procedure results in two datasets from the same system of interest, each with a lower-dimensional latent representation. The cGAN’s task is then to learn a non-linear mapping between latent space representations using a cycle consistency loss (Material and Methods). This procedure constitutes a robust, flexible, and unifying neural network architecture supporting several different integration scenarios, such as between scRNA-seq datasets from different replicates, different library protocols, and different data modalities.

### scAEGAN preserves the cell identity and accurately identifies the cell clusters

To evaluate this concept’s viability and performance, we first tested the scAEGAN by simulated scRNA-seq data using SymSim (X. Zhang et al., 2019). Cells were generated according to a cell population tree, defining several clusters with different distances between them. Using this procedure, two datasets were generated. Each dataset in this simulation had five continuous clusters. In a continuous mode, the cells are positioned along the edges of the tree with a small step size (which is determined by branch lengths and number of cells. Each dataset has 600 cells and 3000 genes simulated with 20 External Variability Factor (EVFs), 12 differential EVFs, and a sigma of 0.4 (Material and Methods, Fig. S1). The number of clusters is preserved in the AE derived low-dimensional embedding. Visual comparison with the translated version of the other domain reveals good agreement (Fig. 1b). We quantified the integration quality by measuring the transfer of labels between the data domains. To this end, we used an SVM to classify cell types in one domain using the cluster assignments. Next, we measured the transferred cell identity agreement with the original identity using the Jaccard Index (JI) and Adjusted Rand Index (ARI) (Material and Methods). The JI calculates 2 by 2 contingency table of agreements and disagreements of the corresponding two vectors of comemberships. Comembership is defined as the pairs of observations that are clustered together. While ARI measures the similarity between the two alternate partitions of the same datasets by the proportion of agreements between the two partitions. The higher the ARI value, the more accurate the clustering, and when the cluster is perfectly matched to the reference criteria, the ARI score equals 1. The scAEGAN preserved the transferred cell identity agreement with the original identity (Fig. 1 c,d).

### scAEGAN integrates datasets across different library protocols

We systematically assessed the ability of scAEGAN-derived feature representations for the integration of different library protocol datasets. To this end we evaluate and compare scAEGAN with Seurat3 as Seurat3 has demonstrated that it can integrate two datasets using different library protocols. It has performed better than Liger(Welch et al., 2019) and scMerge(Lin et al., 2019) when integrating datasets across different single-cell RNA sequencing protocols (Mereu et al., 2020). We evaluated and compared scAEGAN with Seurat 3 using an easier translation task by using ARI and JI as evaluation metrics. We first analyzed the case where we have two versions of the same protocol (CelSeq to CelSeq2) and contrasted this with the more challenging task of integrating two different protocols, e.g., fluidigm F1 with CelSeq. Seurat3 performed well on the easy task (0.62 ARI, 0.52 JI, Fig 3e). Yet, scAEGAN outperformed Seurat 3 in this task (0.88 ARI, 0.82 JI, Fig 3a, b, e). Interestingly, the concatenated architecture (0.38 ARI, 0.32 JI, Fig. 3c, 3d, e) was outperformed by Seurat3. Notably, even the cGAN outperformed Seurat3 (Fig 3e). For the more challenging task (fluidigm F1 with CelSeq), while the performance of scAEGAN dropped (0.66 ARI, 0.62 JI), it still outperformed all other methods and architectures. We observed that even with this challenging task scAEGAN obtained finer granularity in terms of added value to clustering (Fig.3b). We noted that the concatenated and cGAN outperformed Seurat3 in this task. Similar results were obtained in integrating Celseq2 and SMARTseq (Fig. S1). scAEGAN outperformed (0.78 ARI, 0.69 JI) all other methods and architectures. We also evaluated the scAEGAN’s robustness by reducing the no of cells by randomly selecting a % of cells (20, 40, 60, and 80) and computing the ARI for each case. We observed that reducing the number of cells diminishes the performance of Seurat3 and AE-concatenated. Interestingly, when reducing the number of cells, scAEGAN outperforms Seurat3 and AE-concatenated (Fig. S2) significantly, thus suggesting the better robustness of scAEGAN compared to Seurat3 and AE-concatenated.

To assess the GAN cycle consistency loss contribution, we fused the two latent representations by concatenation instead of learning a mapping (Material and Methods). This caused a dramatic drop in performance (0.93 to 0.45 ARI, 0.89 to 0.40 JI, Fig. 2a). Notably, this significant drop occurred despite the simplified situation of well-separated simulated clusters where the latent space’s dimensionality was the same for the two data domains. The analysis using simulated data was repeated for several different cases; all results were in accordance with the above observations (Material and Methods, Fig. S1). Therefore, we concluded that the proposed architecture is sufficient to perform the integration. Furthermore, the analysis also demonstrated the importance of learning a non-linear relationship between the two latent spaces.

**Figure 2.**
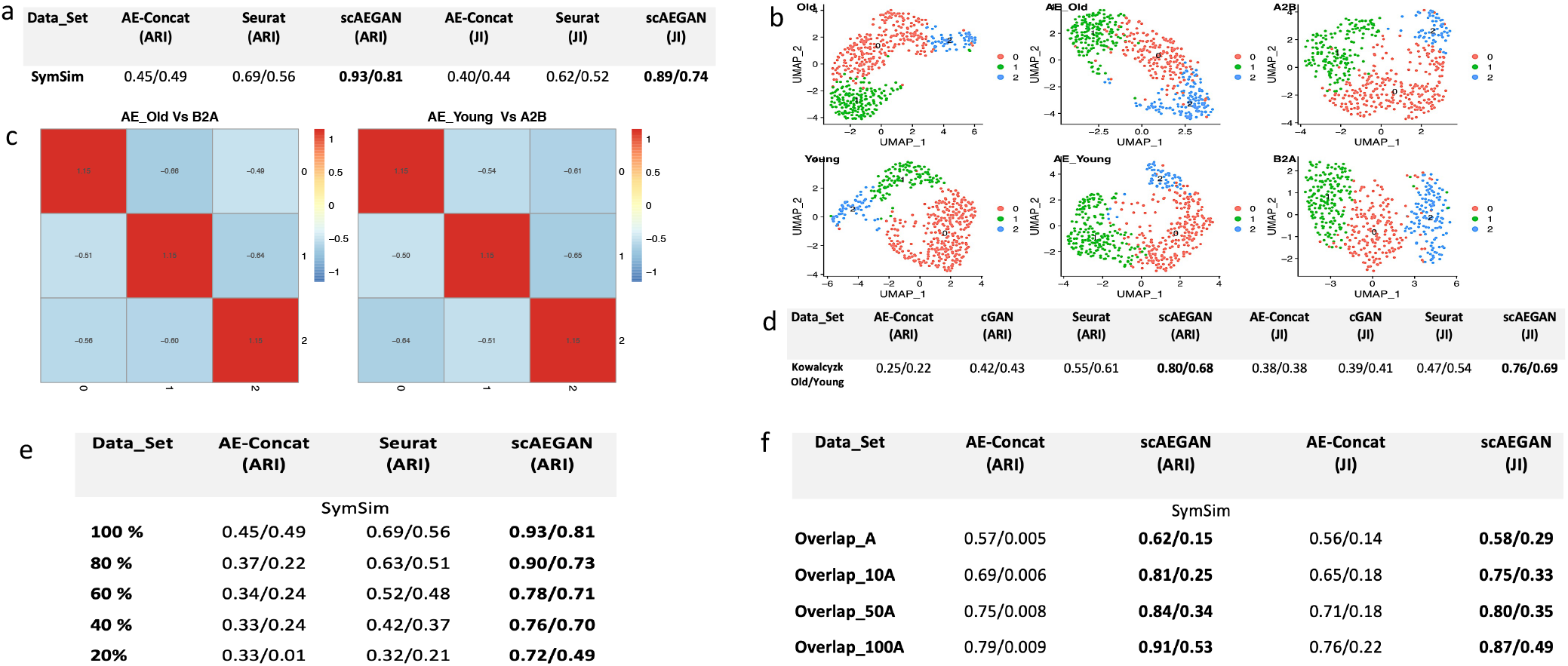
scAEGAN shows robust performance, while integrating datasets from the same platforms, **a)** ARI and JI values depicting the outperformance of scAEGAN with AE-concatenated and Seurat, **b,c,d)** scaAEGAN outperforms other methods for integrating a real scRNA-seq SMARTseq dataset from two mouse strains (Old and Young), **e)** scAEGAN performs better, even the certain percentage of cells are removed from two datasets, **f)** scAEGAN shows robust performance, when there is an imbalance of cell types in two datasets.

**Figure. 3.**
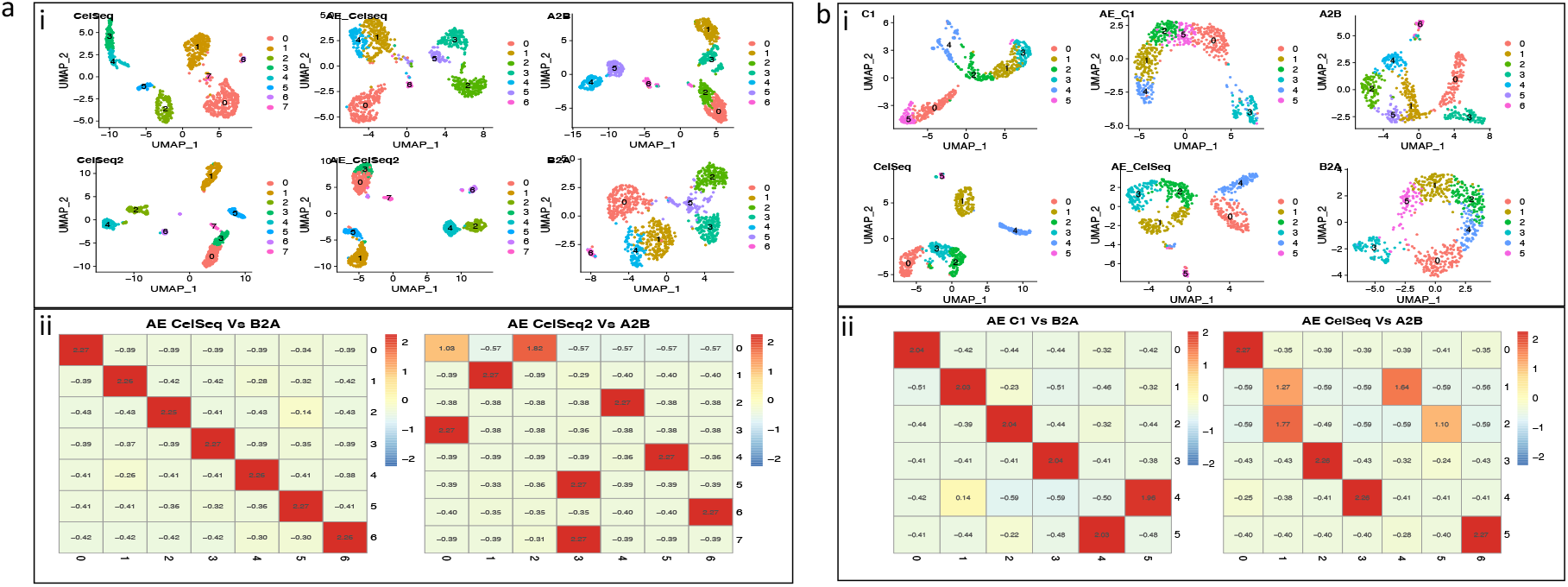
**a,b** Integration results of scAEGAN with across platforms data (CelSeq, CelSeq2, Fluidigm C1) **a,b)** scAEGAN results show better translation of the domains, while maintaining the cluster granularity in the respective domains, while integrating the datasets from CelSeq, CelSeq2 and Fluidigm C1.

**Figure. 3.**
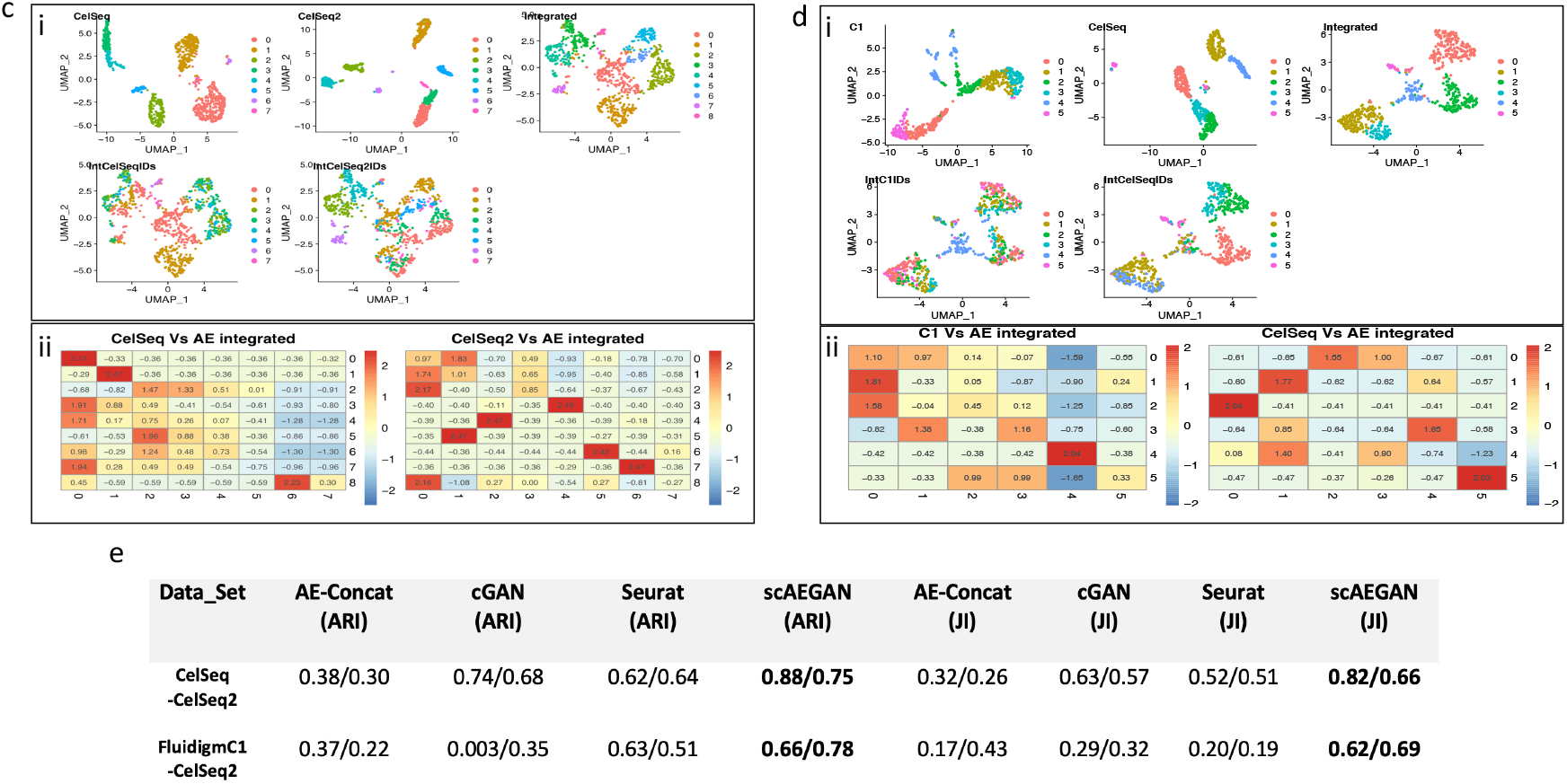
**c,d,e** Integration results of AE-Concatenated with across platforms data (CelSeq, CelSeq2, Fluidigm C1) and its quantitative comparison with scAEGAN, **c, d)** The results from the AE-Concatenated shows its bad performance while integrating the datasets from CelSeq, CelSeq2 and Fluidigm C1, **e)** scAEGAN results shows its outperformance as compared to AE-Concatenated, Seurat and cGAN for integrating data across different platforms.

Next, we asked whether we could learn integration of the input datasets using a cycleGAN without employing an Autoencoder first to project the data into a latent space. This would conceptually correspond to a pixel-by-pixel translation between images. The cycleGAN showed better performance on the simulated datasets (0.99 ARI, 0.92 JI). But when using a real scRNA-seq SMARTseq dataset from two mouse strains (Johansen & Quon, 2019), a reduced performance compared to scAEGAN (0.80 to 0.42 ARI, 0.76 to 0.39 JI, Fig 2b,c,d). Interestingly, the dataset contains several less-informative PCA components likely representing noise in the original data, making it difficult to learn a stable non-linear mapping between the two domains (Fig.S2). The effect of AE training on the two mouse strains datasets retains the most informative PCA components. It removes the components with noise, thus facilitating a linear stable mapping between the two domains (Fig.S2). We also evaluated the robustness of scAEGAN in a simulated setting when we had an imbalance of cell types in two datasets. The imbalance setting ranges from having one dataset with no cluster to removing 10, 50, and 100 cell types from that cluster. scAEGAN performed well (0.62 ARI, 0.58 JI) compared to other methods and architectures (Fig 2f). To further evaluate the robustness of the scAEGAN, we reduced the no of cells by randomly selecting the % of cells (20,40,60, and 80) in the simulated dataset. We observed that scAEGAN outperforms Seurat3 (Fig 2e), thus suggesting the robustness of scAEGAN as compared to Seurat3. Finally, we compared our analysis of the simulated data and the mouse dataset with Seurat3. Overall, the scAEGAN was more successful than Seurat 3 in transferring the labels correctly, whereas Seurat3 was better than the concatenated architecture, thus further supporting the importance of cycleGAN learning.

### scAEGAN outperforms existing methods for the integration of paired and unpaired multi-omic datasets

Aiming for generality, we investigated the integration of multi-omic datasets. To this end, we conducted the integration of scRNA-seq and scATAC-seq data as a case study. When the scRNA-seq and scATAC-seq data are collected from different cells, here referred to as unpaired data, it also includes the challenge of having different samples. In the paired case, both data-modalities are collected from the same cell. Thus, integration of scRNA-seq with scATAC-seq data could either be of paired or unpaired nature. Recent progress has mainly targeted the unpaired case. Tools such as Seurat3 and MOFA+ have demonstrated promising results. A recent upgrade, Seurat4, is the first attempt to our knowledge targeting the paired data-integration challenge. We evaluated the architectures using both paired (Fig 4 a,b,c) and unpaired data (Fig. 4 d,e). As for the previous settings using the Jaccard Index and Adjusted Rand Index as quality measures for quantifying the integration quality, scAEGAN outperforms Seurat 3 and Seurat 4, even when discarding the pairing information between the two modalities (Fig. 4d). To further assess the robustness of scAEGAN, we evaluated the performance of scAEGAN, by removing the % of cells in both paired data and unpaired data. We observed that the performance of Seurat4 decreases with the decrease in the number of cells compared to the scAEGAN. The scAEGAN outperforms Seurat4 (Fig. 4e), thus suggesting better robustness than the Seurat4.

**Figure. 4.**
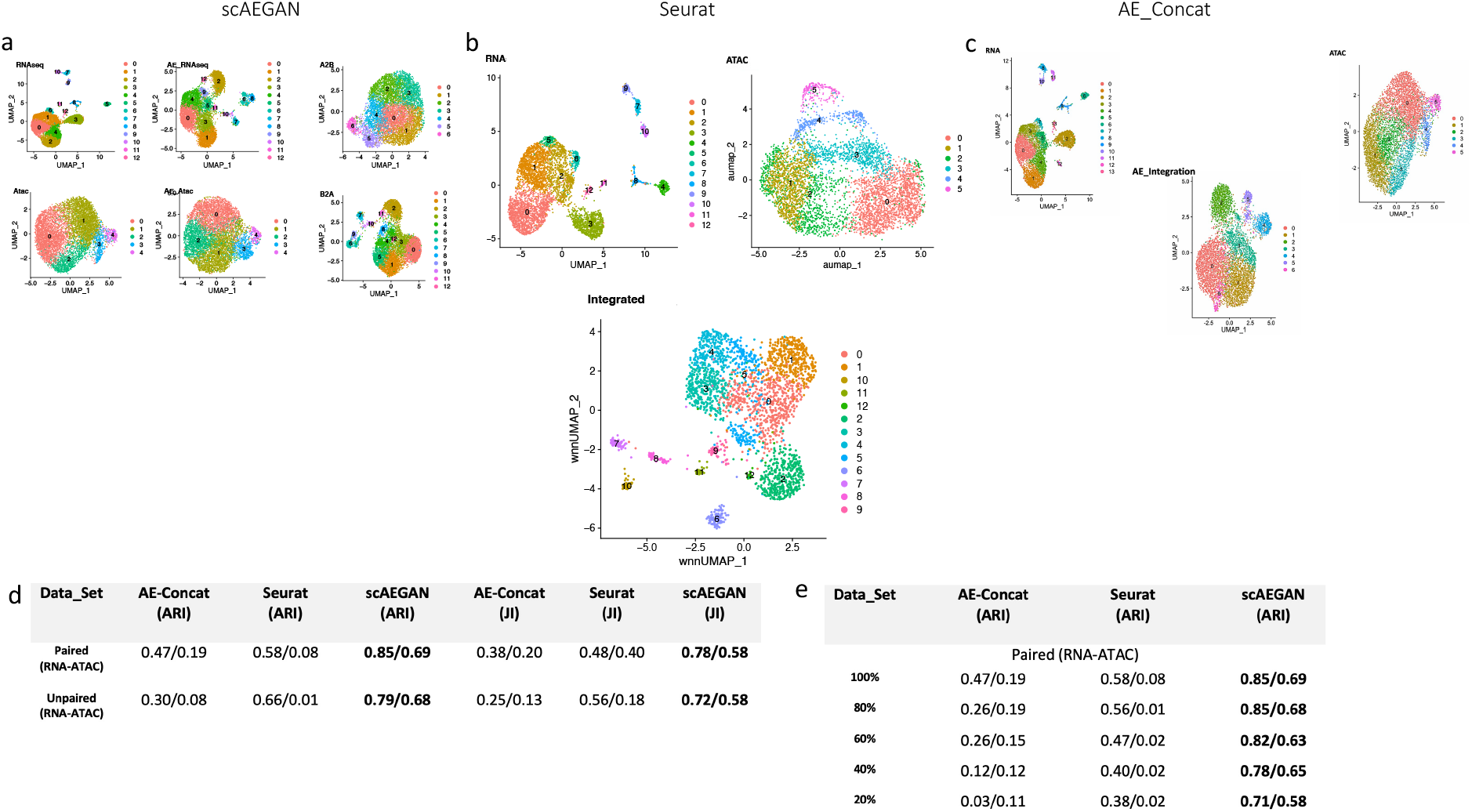
Multimodal integration results of scAEGAN with paired scRNA-seq and scATAC-seq data. The unpaired case is simulated by randomizing the pairing information, **a,b,c,d)** scAEGAN outperforms AE-Concatenated and Seurat 4, even when discarding the pairing information between the two modalities **e)** scAEGAN shows robust performance, even when certain % of cells are removed from each modality.

### scAEGAN facilitates predicting one modality from another modality

To further investigate the efficacy of scAEGAN we attempted to predict one modality from another modality. We trained the scAEGAN on scRNA-seq and tried to predict the scATAC-seq. scAEGAN outperforms Babel, where scAEGAN achieved a Pearson correlation (0.60) as compared to Babel (0.55).

### Concluding remarks

Here we describe scAEGAN, a unifying end-to-end unsupervised single-cell data integration method combining an AE architecture for efficient representation of scRNA-seq data with a CycleGAN network for translation across datasets. We demonstrate the sufficiency in the sense that such a unifying machine learning architecture is able to achieve state-of-the-art or better performance by tackling seemingly “different” integration challenges. Anchoring-based methods, such as Seurat, have a strong domain of applicability and performance when the different datasets are “close” or “similar” to each other. This is natural since the method is predicated on the assumption of “shared” anchors. Yet, when the datasets are either too dissimilar or when there is a need to perform predictions out-of-the-sample, the anchoring approach is limited. For example, as for the challenge of predicting scATAC data from scRNA, machine learning techniques such as Babel are superior to the anchoring approach. Yet, thus far, machine learning methods such as Babel have not yet been able to reach the performance of Seurat on a task such as clustering and integration of unpaired omics data. Here we find that scAEGAN is on par and better than the anchoring technique when different datasets are similar while still being beyond state-of-the-art on prediction tasks, i.e., beating Babel. Our evaluations using the concatenated AE support the interpretation that the critical reason for our success hinges upon that the AE respects each sample’s uniqueness and protocol, whereas the cGAN exploits the similarity in the data distributions in the latent space. Thus, we do not require similarity in the original dataspace, and we can learn a mapping in the latent space across different conditions, thus enabling a predictive capacity. It is an open and interesting problem whether our concept of representation and mapping can be implemented in a different and potentially even more effective manner, thus setting the stage for further development of unifying data integrative frameworks. In summary, the feasibility and robustness of an integrated embedding and learning architecture suggest the broad applicability of such an architecture for integrating data from different replicates, treatments, species, tissues, or sequencing platforms.

## DATA AVAILABILITY

All data and software are publicly accessible, as detailed in the Materials & Methods section. The code is available at https://github.com/sumeer1/scAEGAN.

## SUPPLEMENTARY DATA

Supplementary Data are available at NAR online.

## REFERENCES

Argelaguet, R., Arnol, D., Bredikhin, D., Deloro, Y., Velten, B., Marioni, J. C., & Stegle, O. (2020). MOFA+: A statistical framework for comprehensive integration of multi-modal single-cell data. Genome Biology, 21(1), 111. https://doi.org/10.1186/s13059-020-02015-1

Arjovsky, M., Chintala, S., & Bottou, L. (2017). Wasserstein Generative Adversarial Networks. https://doi.org/10.5555/3305381

Chen, S., Lake, B. B., & Zhang, K. (2019). High-throughput sequencing of the transcriptome and chromatin accessibility in the same cell. Nature Biotechnology 2019 37:12, 37(12), 1452–1457. https://doi.org/10.1038/s41587-019-0290-0

Eraslan, G., Simon, L. M., Mircea, M., Mueller, N. S., & Theis, F. J. (2019). Single-cell RNA-seq denoising using a deep count autoencoder. Nature Communications, 10(1), 1–14. https://doi.org/10.1038/s41467-018-07931-2

Haghverdi, L., Lun, A. T. L., Morgan, M. D., & Marioni, J. C. (2018). Batch effects in single-cell RNA-sequencing data are corrected by matching mutual nearest neighbors. Nature Biotechnology, 36(5), 421–427. https://doi.org/10.1038/nbt.4091

Hao, Y., Hao, S., Andersen-Nissen, E., Mauck, W. M., Zheng, S., Butler, A., Lee, M. J., Wilk, A. J., Darby, C., Zager, M., Hoffman, P., Stoeckius, M., Papalexi, E., Mimitou, E. P., Jain, J., Srivastava, A., Stuart, T., Fleming, L. M., Yeung, B., … Satija, R. (2021). Integrated analysis of multimodal single-cell data. Cell. https://doi.org/10.1016/j.cell.2021.04.048

Hinton, G. E., & Salakhutdinov, R. R. (2006). Reducing the dimensionality of data with neural networks. Science, 313(5786), 504–507. https://doi.org/10.1126/science.1127647

Johansen, N., & Quon, G. (2019). ScAlign: A tool for alignment, integration, and rare cell identification from scRNA-seq data. Genome Biology, 20(1), 166. https://doi.org/10.1186/s13059-019-1766-4

Kingma, D. P., & Ba, J. L. (2015, December 22). Adam: A method for stochastic optimization. 3rd International Conference on Learning Representations, ICLR 2015 - Conference Track Proceedings. https://arxiv.org/abs/1412.6980v9

Korsunsky, I., Millard, N., Fan, J., Slowikowski, K., Zhang, F., Wei, K., Baglaenko, Y., Brenner, M., Loh, P. ru, & Raychaudhuri, S. (2019). Fast, sensitive and accurate integration of single-cell data with Harmony. Nature Methods, 16(12), 1289–1296. https://doi.org/10.1038/s41592-019-0619-0

Li, G., Fu, S., Wang, S., Zhu, C., Duan, B., Tang, C., Chen, X., Chuai, G., Wang, P., & Liu, Q. (2022). A deep generative model for multi-view profiling of single-cell RNA-seq and ATAC-seq data. Genome Biology, 23(1), 20. https://doi.org/10.1186/s13059-021-02595-6

Lin, Y., Ghazanfar, S., Wang, K. Y. X., Gagnon-Bartsch, J. A., Lo, K. K., Su, X., Han, Z. G., Ormerod, J. T., Speed, T. P., Yang, P., & Yang, J. Y. H. (2019). ScMerge leverages factor analysis, stable expression, and pseudoreplication to merge multiple single-cell RNA-seq datasets. Proceedings of the National Academy of Sciences of the United States of America, 116(20), 9775–9784. https://doi.org/10.1073/pnas.1820006116

Maas, A. L., Hannun, A. Y., & Ng, A. Y. (2013). Rectifier Nonlinearities Improve Neural Network Acoustic Models.

Mereu, E., Lafzi, A., Moutinho, C., Ziegenhain, C., McCarthy, D. J., Álvarez-Varela, A., Batlle, E., Sagar, Grün D., Lau, J. K., Boutet, S. C., Sanada, C., Ooi, A., Jones, R. C., Kaihara, K., Brampton, C., Talaga, Y., Sasagawa, Y., Tanaka, K., … Heyn, H. (2020). Benchmarking single-cell RNA-sequencing protocols for cell atlas projects. Nature Biotechnology, 38(6), 747–755. https://doi.org/10.1038/s41587-020-0469-4

Qin, Y., Mitra, N., & Wonka, P. (2018). How does Lipschitz Regularization Influence GAN Training? Lecture Notes in Computer Science (Including Subseries Lecture Notes in Artificial Intelligence and Lecture Notes in Bioinformatics), 12361 LNCS, 310–326. http://arxiv.org/abs/1811.09567

Shafer, M. E. R. (2019). Cross-Species Analysis of Single-Cell Transcriptomic Data. In Frontiers in Cell and Developmental Biology (vol. 7, p. 175). Frontiers Media S.A. https://doi.org/10.3389/fcell.2019.00175

Stuart, T., Butler, A., Hoffman, P., Hafemeister, C., Papalexi, E., Mauck, W. M., Hao, Y., Stoeckius, M., Smibert, P., & Satija, R. (2019). Comprehensive Integration of Single-Cell Data. Cell, 177(7), 1888–1902.e21. https://doi.org/10.1016/j.cell.2019.05.031

Stuart, T., & Satija, R. (2019). Integrative single-cell analysis. In Nature Reviews Genetics (vol. 20, Issue 5, pp. 257–272). Nature Publishing Group. https://doi.org/10.1038/s41576-019-0093-7

Tran, H. T. N., Ang, K. S., Chevrier, M., Zhang, X., Lee, N. Y. S., Goh, M., & Chen, J. (2020). A benchmark of batch-effect correction methods for single-cell RNA sequencing data. Genome Biology, 21(1), 12. https://doi.org/10.1186/s13059-019-1850-9

Waltman, L., & Van Eck, N. J. (2013). A smart local moving algorithm for large-scale modularity-based community detection. European Physical Journal B, 86(11), 471. https://doi.org/10.1140/epjb/e2013-40829-0

Welch, J. D., Kozareva, V., Ferreira, A., Vanderburg, C., Martin, C., & Macosko, E. Z. (2019). Single-Cell Multi-omic Integration Compares and Contrasts Features of Brain Cell Identity. Cell, 177(7), 1873–1887.e17. https://doi.org/10.1016/j.cell.2019.05.006

Wu, K. E., Yost, K. E., Chang, H. Y., & Zou, J. (2021). BABEL enables cross-modality translation between multiomic profiles at single-cell resolution. Proceedings of the National Academy of Sciences of the United States of America, 118(15). https://doi.org/10.1073/pnas.2023070118

Zhang, X., Xu, C., & Yosef, N. (2019). Simulating multiple faceted variability in single cell RNA sequencing. Nature Communications, 10(1). https://doi.org/10.1038/s41467-019-10500-w

Zhang, Y., & Wang, F. (2021). SSBER: removing batch effect for single-cell RNA sequencing data. BMC Bioinformatics, 22(1). https://doi.org/10.1186/s12859-021-04165-w

Zhu, J. Y., Park, T., Isola, P., & Efros, A. A. (2017). Unpaired Image-to-Image Translation Using Cycle-Consistent Adversarial Networks. Proceedings of the IEEE International Conference on Computer Vision. https://doi.org/10.1109/ICCV.2017.244

